# A new bioinformatic tool to interpret metagenomics / metatranscriptomics results based on the geometry of the clustering network and its differentially gene ontologies (GANGO)

**DOI:** 10.1101/2020.06.10.140103

**Authors:** Antonio Monleon-Getino, Andreu Paytuví-Gallart, Walter Sanseverino, Javier Méndez

## Abstract

High-throughput experimental techniques, such as metagenomics or metatranscriptomics, produce large amounts of data, which interpretation and conversion into understandable knowledge can be challenging and out of reach. We present GANGO, a new algorithm based on the ecological concept of consortium (groups biologically connected) and by using clustering network analysis, gene ontologies and powerful hypothesis test allows the identification and interpretation of complex ecological networks, allowing the identification of the relationship between taxa/genes, the number of groups, their relations and their functionalities using the annotated genes of an organism in a database (e.g. UniProt or Ensembl). Three examples of the use of GANGO are shown: a simulated mixture of fungi and bacteria, alterations in soil fungi communities after a diesel-oil spill and genomic changes in *Saccharomyces cerevisae* due to abiotic stress.

## 1. Background

High-throughput experimental techniques [1], such as metagenomics or metatranscriptomics, produce large amounts of data, which interpretation and conversion into understandable knowledge can be challenging and out of reach.

We present GANGO, a new algorithm that based on the ecological concept of consortium (groups biologically connected) and by using clustering network analysis, gene ontologies and powerful hypothesis test allows the identification and interpretation of complex ecological networks, allowing the identification of the relationship between taxa/genes, the number of groups, their relations and their functionality using the annotated genes of an organism in a database (e.g. UniProt or Ensembl).

Three examples of use of GANGO are shown: a simulated mixture of fungi and bacteria, alterations in soil fungi communities after a diesel-oil spill and genomic changes in *Saccharomyces cerevisae* due to abiotic stress.

## 2. Objectives

Developing a new method (algorithm) in R that allows:

- To identify possible groups of microrganisms (consortia) from metagenomics studies based on a previous function developed in R.
- To provide a biological interpretation of consortia based on different information retrieved from proteins (GO terms, KEGG pathways) annotated in public databases.
- To encapsulate all these functionalities in a function and arrange it in an R library with the aim to allow the scientific community to freely use it.

## 3. Material and methods

In the figure 1 the procedures used by GANGO are shown to obtain a biological interpretation of the described results as well as their basis. GANGO basically starts (input) from a list of OTUs, which are identified during a metagenomics analysis. OTUS are clustered in groups (identified as a consortium), oftentimes, taxonomic or experience-based criteria are used to establish them and observe their meaning.

**Figure 1:**
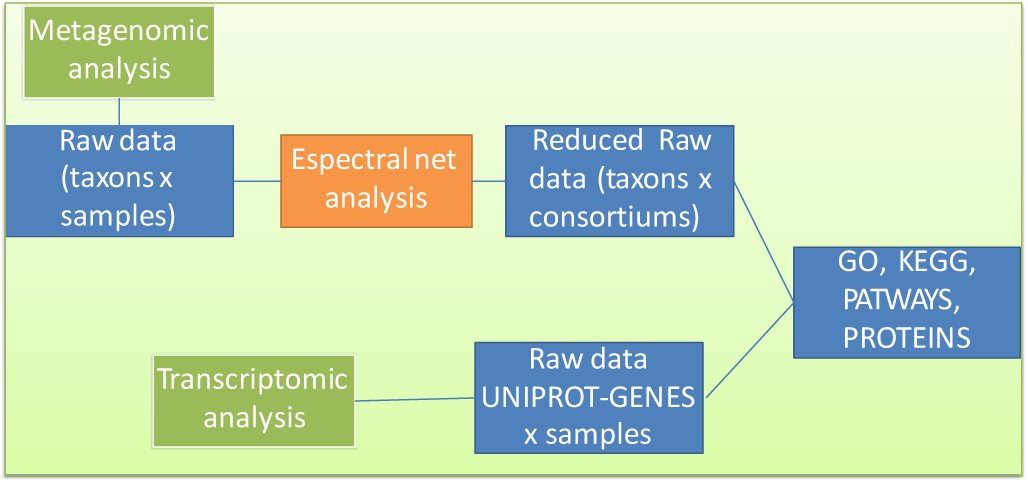
Diagram of the functioning of GANGO

We propose that consortia to be established through network analysis using the **Espectral.CN()** function of the R **library(BDSbiost3)** (Monleón, 2020)[2].

GANGO is able to start from a list of genes, as in Example 3, where it starts from a list of differentially expressed genes from an RNAseq experiment.

As state above, three examples of use of GANGO are shown: (1) a mixture of fungi and bacteria obtained by simulation, (2) alterations in soil fungi communities after a diesel-oil spill obtained in the bibliography [3] and (3) genomic changes in *S. cerevisae* due to abiotic stress obtained during a RNAseq experiment [4][5].

Example 1 was carried out with metagenomics simulated data using the GAIA pipeline [6][7] to identify taxa. This data was *in silico* generated from different genomes at random proportions using wgsim (https://github.com/lh3/wgsim).

Example 2 was carried out with soil fungi taxa (Figure 2) in a soil where fungi communities were studied after a diesel-oil spill, this example was obtained from bibliography. Fungi were identify by biochemical methods (not metagenomic). The diversity data was extracted from: https://www.frontiersin.org/articles/10.3389/fmicb.2017.01862/full [3].

**Figure 2:**
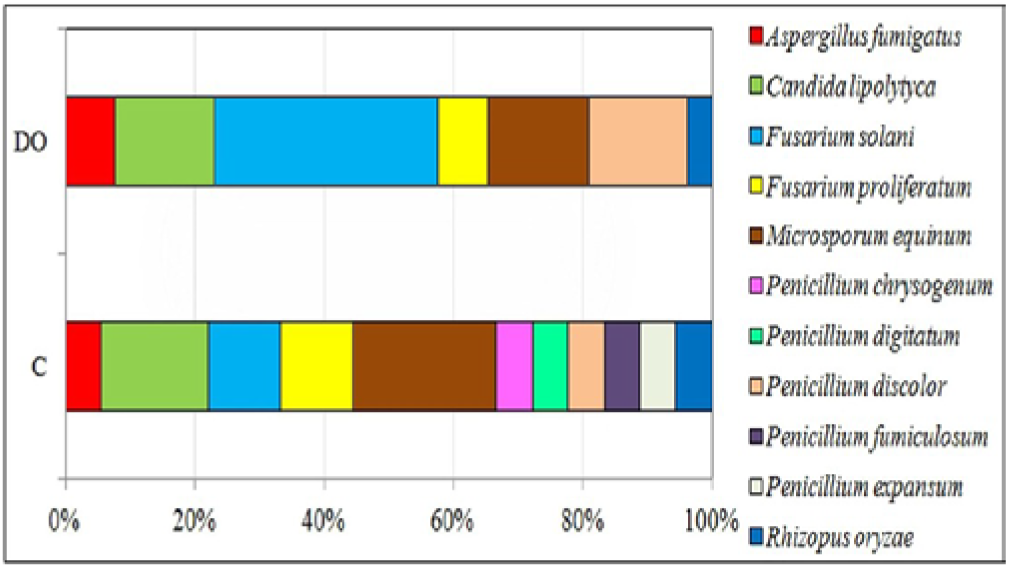
Fungi present differentially in samples soils after diesel-oil spill (C=CONTROL / DO = Diesel OIL). Taxa were obtained from [4].

Example 3 was carried out on *S. cerevisiae* under abiotic stress versus control conditions whose genes were differentially expressed using pipeline AIR [4][5]. All the data and the gene extraction are detailedly explained in Monleón (2020) [8] and comes from a previous training session for the use of AIR by Sequentia Biotech.

**AIR results from this experiment are available at** https://transcriptomics.sequentiabiotech.com/shared/TaskFlow/d11c3523-b381-4778-ab06-d78eddc3a73a/0dd29e9a-111f-4d40-b3ba-b864f308e3d8

## 4. Results

Results are presented in graphical form, and they lies in complex networks of consortia, volcano plots using a Fisher’s exact test of differential proportions, and the use of function **coincidence_analysis()**, which performs Venn diagrams to compare taxa between groups. The function **dif.propOTU.between.groups()**, carries out a statistical test of the proportions between groups. All these functions belong to the library BDbiost3[4].

### First example: microorganism consortia from a simulated metagenomic data

Three consortia from 140 taxa were detected using the geometry analysis of the network formed between taxa data (see figure 4). Networks were obtained using the **Espectral.CN()** function from the R **l**ibrary **BDSbiost3** (Monleón, 2020)[2]. Venn diagram between groups is represented in figure 4 in order to know the composition between groups of microorganisms. GANGO was applied to these consortia and all GO associated were stored for its analysis. The differential GO proportion test, based on the Gene Ontology (GO) terms obtained from UniProt for each group, was computed using the function **dif.propOTU.between.groups()**. Results between groups G2 and G3 are presented in figure 5.

**Figure 3:**
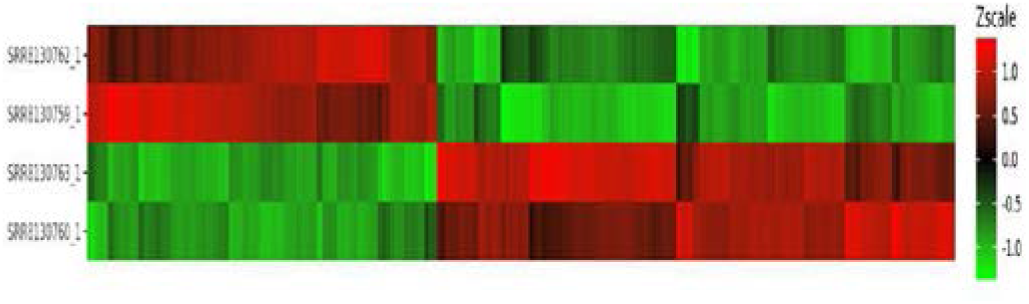
Heatmap for genes and samples differentially expressed of the study of genomic changes in *S. cerevisae* due to abiotic stress obtained during a RNAseq experiment [5].

**Figure 4:**
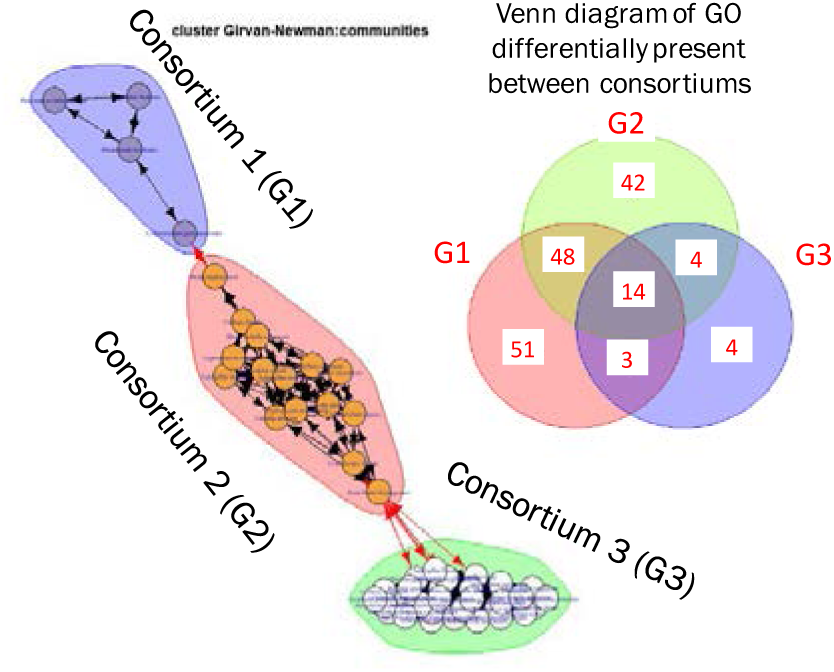
Three microbiological consortia were detected using geometrical analysis of the network formed using Espectral.CN(). Venn diagram of the consortia were represented.

**Figure 5:**
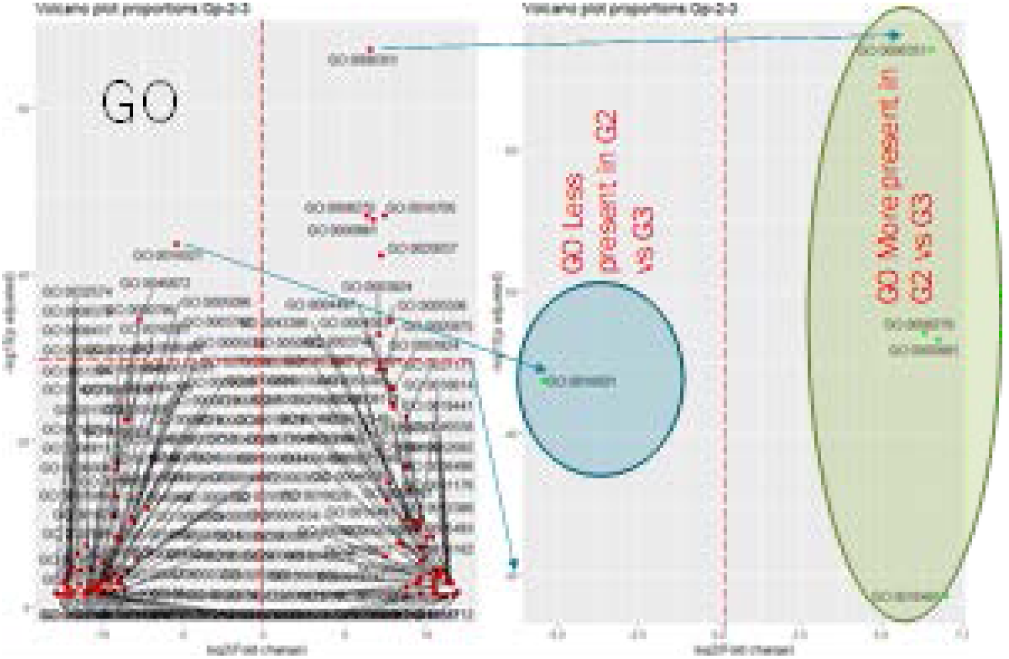
Differential analysis of GO after GANGO algorithm of the consortia G2 and G3, detected previously. Volcano plot (left) where is possible observe GO differentially proportion detected by function dif.propOTU.between.groups(). Detail of the GO differentially proportion analysis (right).

### Second example: GO analysis of soil fungi during diesel-oil spill

In figure 6 it is shown fungi taxa from soil, these communities were studied after a diesel-oil spill [3]. Also, Venn diagram is represented.

**Figure 6:**
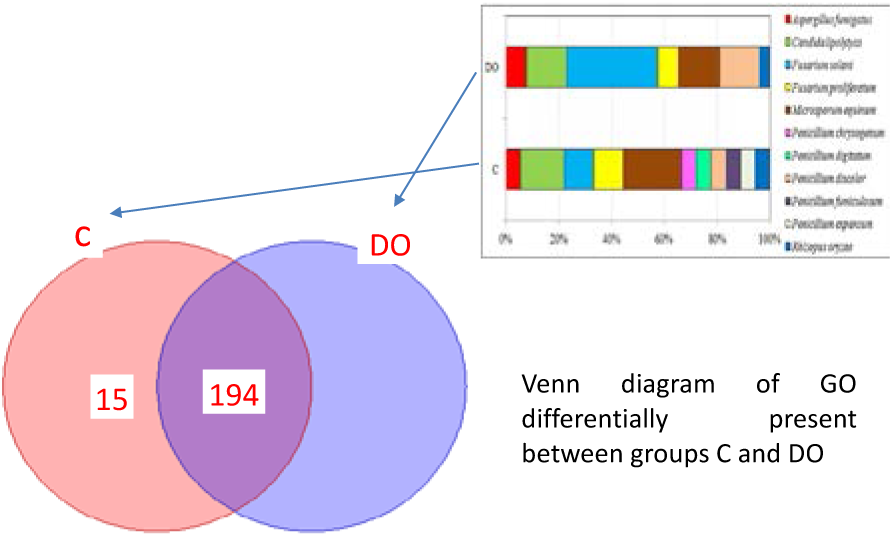
taxa used during the second example. Is possible see fungi taxa from soil, these communities were studied after a diesel-oil spill (C=Control, DO = diesel oil) [4]. Venn diagram is represented.

GANGO was applied to these two groups of taxa (Control group and Diesel oil group) and all GO terms were stored for its analysis. Differential GO proportion test based on the GO terms obtained from UniProt for each group was computed using the function **dif.propOTU.between.groups()**. Statistical tests between groups “Control” and “Diesel oil” are presented in figure 7.

**Figure 7:**
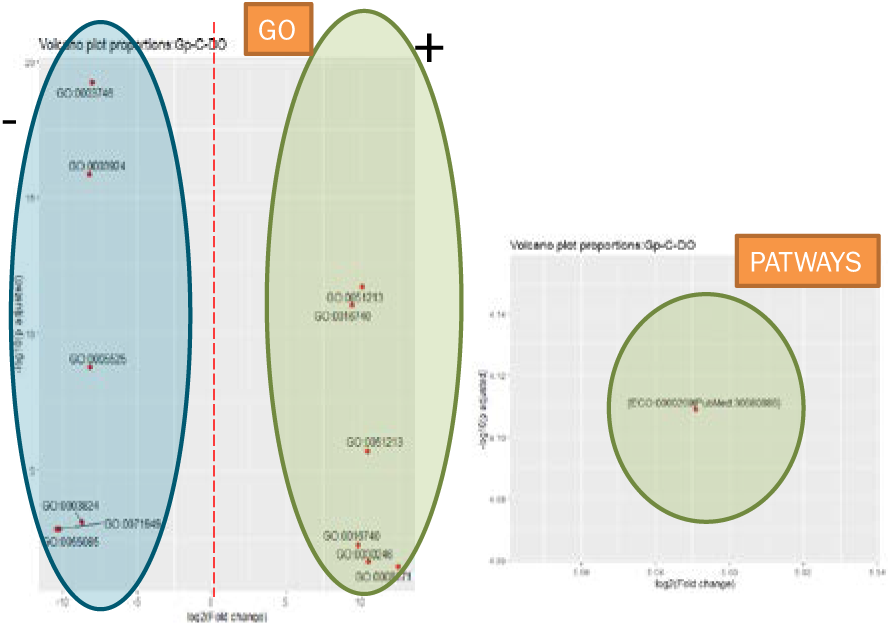
Differential analysis of GO after GANGO algorithm of the taxa of the groups “control” and “diesel oil” of the secons example (fungus in soil). Volcano plot (left) where is possible observe GO differentially proportion detected by function dif.propOTU.between.groups(). Detail of the GO differentially proportion analysis (right).

### Third example: *Saccharomyces cerevisae* differential expression using RNAseq

Differential proportion test based on the GO ontologies obtained from UniProt for each group (analysis of “control” vs “abiotic stress”) is shown in figure 8 as a volcano plot. Where *X* is the fold change and *Y* is the *p*-value (after apply FDR method). The distribution of the dots in the plot (figure 8) can be useful to check if a range of fold-changes is associated with a stronger or a weaker significance of differential expression.

**Figure 8:**
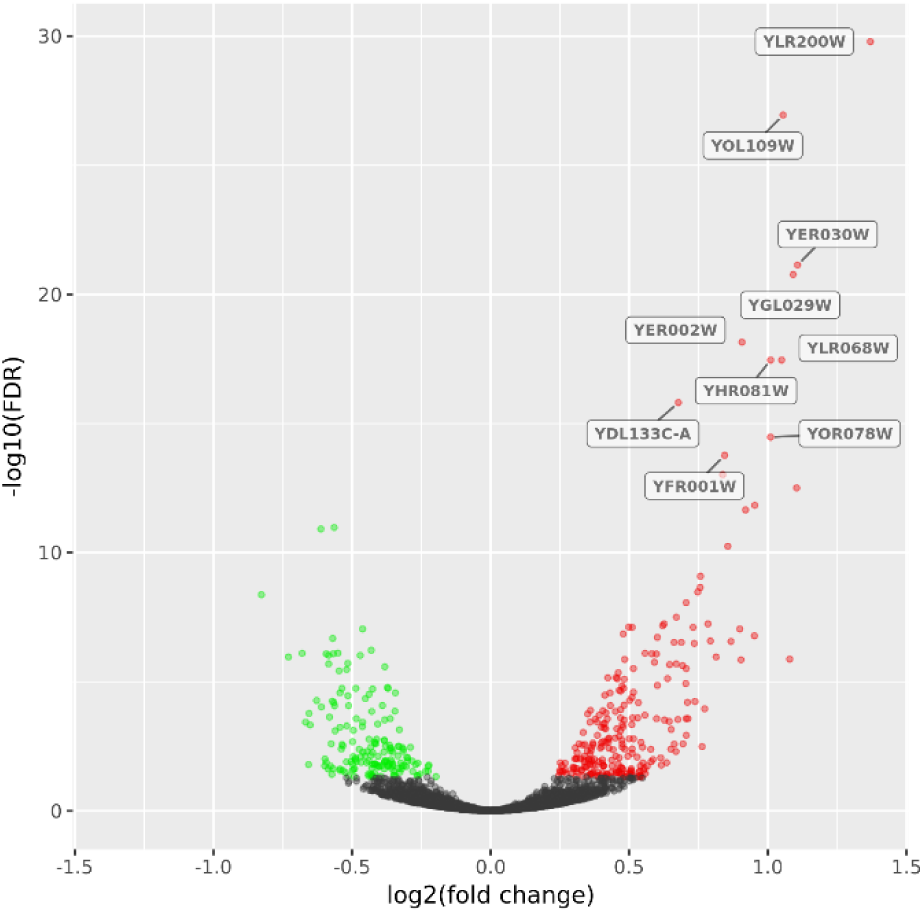
Volcano plot retrieved from the AIR analysis (see section 3. Materials and Methods). In it there are shown differentially expressed genes from *S. cerevisae* in the comparation control vs. stress condition. Black dots represent genes that are not significantly differentially expressed, while red and green dots are the genes that are significantly UP- and DOWN-regulated, respectively.

GANGO was applied to these two groups of differentially expressed genes (genes that are significantly UP- and DOWN-regulated, see figure 8) and all GO terms associated were stored for their analysis. Differential GO proportion test based on the GO terms obtained from UniProt for each group was computed using the function **dif.propOTU.between.groups()**. Statistical tests between groups “control” and “abiotic stress” are presented in figure 9.

**Figure 9:**
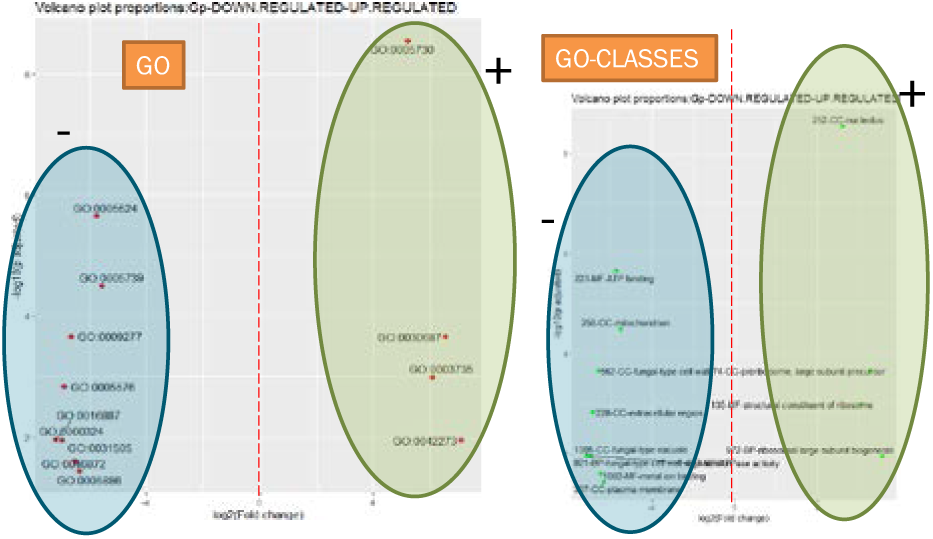
Differential analysis of GO terms after GANGO algorithm from the genes differentially expressed. Differentially proportion anlysis of the GO terms (left) and GO classes/names (right). Circles show differential GO terms from UP- and DOWN-regulated genes detected between the experimental groups.

## GANGO algorithm

GANGO algorithm can be found in the GANGO package for R on Github: https://github.com/amonleong/GANGO

**Function GANGO description**

GANGO(

taxon,

group,

callto_old_file = F,

Gene_yes = F,

Gene_list,

All_fields = F)

- Taxon: taxa identified in a metagenomic analysis
- Group: experimental groups
- callto_old_file: reuse an old file
- Gene_yes: use a list of genes (T/F)
- Gene_list: list if genes
- All_fields: obtain all GO files

## 5. Conclusions and comments

GANGO is an R package to perform gene ontology analyses on consortia groups (either genes or taxa). We strongly believe that it is an innovative alternative for researchers when seeking out microbiological consortia, considering the biological interpretation using genes annotated in UNIPROT and extracting the gene ontologies associated with them.

In addition, GANGO might be a useful implementation for diagnostic tools or in determining feasible microorganisms or derived products to formulate products such as fertilizers, prebiotics or probiotics products. In this context, GANGO is a perfect complement of current gene ontology analyses tto make the interpretation of the results more understandable.

## ANNEX: EXAMPLE OF LIBRARY GANGO

Download GANGO at: https://github.com/amonleong/GANGO

#First example: GENES ANOTATED in Diversity of Fungi in Soil Polluted with Diesel Oil

\# DATA FROM:

https://www.frontiersin.org/articles/10.3389/fmicb.2017.01862/full #Taxa identified in the experiment COMPLETE taxons_dieseloil_control<-c(“Aspergillus fumigatus”, “Candida lipolytica”, “Fusarium solani”, “Fusarium proliferatum”, “Microsporum equinum”, “Penicillium discolor”, “Rhizopus oryzae”, “Aspergillus fumigatus”, “Candida lipolytica”, “Fusarium solani”, “Fusarium proliferatum”, “Microsporum equinum”, “Penicillium chrysogenum”, “Penicillium digitatum”, “Penicillium discolor”, “Penicillium discolor”,”Penicillium fumiculosum”, “Penicillium expansum”, “Rhizopus oryzae”)

#Experimental groups (C=ontrol, DO=diesel oil) group_dieseloil_control<-c(“DO”,”DO”,”DO”,”DO”,”DO”,”DO”,”DO”, “C”,”C”,”C”,”C”,”C”,”C”,”C”,”C”,”C”,”C”,”C”,”C”)

#Taxa identified in the experiment SIMPLIFICATION taxons_dieseloil_control1<-c(“Aspergillus fumigatus”, “Candida lipolytica”, “Penicillium expansum”, “Rhizopus oryzae”)

#Experimental groups (C=ontrol, DO=diesel oil) group_dieseloil_control1<-c(“DO”,”DO”, “C”,”C”) #Use GANGO to search anotated GO, PATWAY, KEGG,

PROTEINS(Map GO from taxa) in UNIPROT #Save results in: setwd(“C:/Users/Toni/Downloads”)

#Map GO from UNIPROT using GANGO (Maybe a lot of time!!!!)

GO.detaxon<-GANGO(taxon=taxons_dieseloil_control1, group=group_dieseloil_control1,callto_old_file = F,All_fields = F,Gene_yes = F)

\# Map GO from UNIPROT using GANGO (Maybe a lot of time!!!!)

#GO.detaxon1<-GANGO(taxon=taxons_dieseloil_control1, group=group_dieseloil_control1,callto_old_file = F,All_fields = T,Gene_yes = F)

#GANGO results: cross_table<-GO.detaxon[[1]] #Cross table of GO and groups res3<-GO.detaxon[[3]]

#Results of differentially proportions using dif.propOTU.between.groups() of library(BDSbiost3) library(BDSbiost3) #Download at https://github.com/amonleong/BDSbiost3 dif.propOTU.between.groups(matriu = cross_table, vector.labels=colnames(cross_table), alfa=0.05, lim.padj = 0.00001, lim.log2FoldChange =9)

#coincidence analysis between groups using library(BDSbiost3) coincidence.analysis(cross_table)

